# Ketamine Modulates Endogenous Opioid Peptide Signaling and Reduces Heroin-Seeking in Male Long-Evans Rats

**DOI:** 10.64898/2026.09.01.748554

**Authors:** Adam M. Dawoud, Achla Gupta, Ivone Gomes, Dylan J. Baker, Julia E. Shapiro, Lakshmi Devi, Yasmin L. Hurd, Aya Osman

**Affiliations:** Addiction Institute of Mount Sinai, Icahn School of Medicine at Mount Sinai, New York, New York, USA; Department of Neuroscience, Friedman Brain Institute, Icahn School of Medicine at Mount Sinai, New York, New York, USA; Department of Pharmacological Sciences, Icahn School of Medicine at Mount Sinai, New York, New York, USA; Department of Psychiatry, Friedman Brain Institute, Icahn School of Medicine at Mount Sinai, New York, New York, USA; Department of Biological Sciences, University at Albany – State University of New York, Albany, New York, USA

## Abstract

Opioid Use Disorder (OUD) continues to be a pressing public health crisis, marked by high relapse rates and limited treatment options. Ketamine, a non-competitive N-Methyl-D-Aspartate receptor (NMDAR) antagonist, has emerged as a promising therapeutic for numerous psychiatric illnesses due to its robust antidepressant properties and glutamate-mediated synaptic plasticity. However, ketamine’s therapeutic mechanisms remain elusive, with emerging evidence implicating the endogenous opioid system. Furthermore, few studies have investigated ketamine’s potential as an intervention for OUD. Accordingly, this study assessed ketamine’s ability to rewire endogenous opioid and glutamatergic circuitry and reduce heroin-seeking activity in male Long-Evans rats. Using a multi-cohort design, we investigated ketamine’s effects on opioid-related transcriptional, protein, and signaling properties in key brain regions associated with reward and addiction, and subsequently on heroin-seeking using a self-administration paradigm. To investigate mechanisms of plasticity, we also quantified transcriptional changes to NMDAR subunits. Molecular analyses demonstrated increased expression of endogenous opioid system-related genes and endogenous opioid peptides at the protein level, elevated mu-opioid receptor protein levels, and enhanced G protein activity, suggesting increased receptor function. Notably, transcriptional changes were particularly prominent in the shell subregion of the nucleus accumbens (NAc). We also revealed that ketamine altered NMDAR subunit expression in the medial prefrontal cortex and NAc, indicating circuit-level synaptic remodeling. Behavioral analyses revealed that ketamine significantly reduced heroin-primed, but not cue-induced, seeking activity. Together, these findings suggest ketamine reduces acute relapse vulnerability to opioids through coordinated modulation of opioid and glutamatergic systems, highlighting its potential as a novel intervention for OUD.

## Introduction

The negative implications of Opioid Use Disorder (OUD) have long daunted our society. In 2020, over 16 million people globally suffered from OUD, with more than 120,000 fatal opioid-related overdoses [1]. Moreover, rates of opioid-related overdoses in the United States have increased nearly five-fold from 2015 to 2020 [2], largely attributed to the influx of synthetic opioids into the illicit drug market. Current treatments for OUD consist of opioid agonist therapy with buprenorphine or methadone. Given the increasing rates of OUD and the limited effective treatment options, there is a pressing need to identify safer and more effective therapeutics.

Ketamine is a non-competitive antagonist of the N-Methyl-D-Aspartate receptor (NMDAR) with rapid and long-lasting antidepressant effects following administration of a single dose [3]. One of its proposed antidepressant mechanisms involves antagonism of NMDARs on GABAergic interneurons in the prefrontal cortex (PFc). This disinhibits glutamatergic projections to the nucleus accumbens (NAc), increasing brain-derived neurotrophic factor release and synaptogenesis through the mechanistic target of rapamycin pathway, in the PFc-NAc circuit [4]. Although ketamine research has largely focused on NMDAR mechanisms, growing evidence implicates the endogenous opioid system in its therapeutic effects.

In 2018, administration of naltrexone, a mu-opioid receptor (MOR) antagonist, in conjunction with ketamine was shown to abolish ketamine’s antidepressant effects [5]. This study sparked further investigation into ketamine’s endogenous opioid system mechanisms [6, 28]. Indeed, ketamine has been shown to bind MOR and kappa opioid receptors [7]. A recent cryo-EM study reported that ketamine directly engages MORs [8]. Similarly, transcription of Oprm1, the gene encoding MOR, is increased in rats one hour following 10 mg/kg intraperitoneal (ip) ketamine treatment [9]. Additionally, ketamine potentiates opioid-induced extracellular regulated kinase (ERK1/2) phosphorylation in a cell line exclusively expressing MORs [10, 13]. Finally, ketamine has been shown to potentiate endogenous opioid peptide-induced G-protein activation, suggesting that it acts as a positive allosteric modulator (PAM) of opioid receptors [10]. Ketamine metabolites lacking NMDAR activity also function as opioid receptor PAMs [10]. While research investigating ketamine’s potential to alleviate symptoms of OUD is limited, these pharmacological features suggest it may be an effective intervention for OUD patients suffering from a dysregulated endogenous opioid system.

Clinically, high-dose intramuscular ketamine (2.0 mg/kg) during psychotherapy sessions significantly increased two-year abstinence rates in detoxified heroin-dependent patients compared to lower-dose controls [14]. A follow-up study demonstrated that multiple sessions further improved one-year abstinence outcomes [15]. Preclinically, a single 10 mg/kg ip ketamine injection was sufficient to reverse morphine-conditioned place preference in mice [16], with similar findings in rats [17].

While ketamine has consistently shown promise as a therapeutic for OUD, no studies have specifically assessed ketamine’s potential to mitigate symptoms of opioid relapse. Furthermore, the mechanisms by which ketamine elicits its therapeutic effects remain unclear. Here, we assessed dose-, time-, and brain region-dependent effects of acute ketamine treatment on opioid and glutamatergic signaling. Subsequently, we tested ketamine’s effects on heroin seeking using a rodent self-administration model.

## Methods

### Animals and Housing

All experiments were conducted using adult male Long-Evans rats (Charles River, Wilmington, MA) housed in temperature-controlled environments under a 12-hour light/dark cycle, with ad libitum access to food and water. The experiments adhered to the National Institutes of Health guidelines and received approval from the Institutional Animal Care and Use Committee (IACUC) at the Icahn School of Medicine at Mount Sinai. Animals were housed in Association for Assessment and Accreditation of Laboratory Animal Care (AALAC)-accredited facilities.

### Acute Ketamine Cohorts

Three acute cohorts were utilized. Ketamine hydrochloride (Ketasthesia, racemic, Henry Schein) was diluted in 0.9% saline to a 100 mg/mL stock solution. In Cohort 1, rats received intraperitoneal injections of either saline (n=4), 3 mg/kg ketamine (n=12), or 10 mg/kg ketamine (n=12). Saline treated animals were sacrificed at 24 hours, and 4 animals from each ketamine dose group were sacrificed at 24 hours (n=8), 72 hours (n=8), and 168 hours (n=8). In Cohort 2, the 3 mg/kg treatment was eliminated based on Cohort 1 findings and sample sizes were increased: saline (n=8) and 10 mg/kg ketamine (n=24). Saline animals were sacrificed at 24 hours, and ketamine animals were sacrificed at 24 hours (n=8), 72 hours (n=8), and 168 hours (n=8). In Cohort 3, the 24- and 72-hour timepoints were eliminated for the 10 mg/kg ketamine group; instead, rats were sacrificed 168 hours (n=8) after treatment. Saline-treated animals (n=8) were sacrificed 24 hours after treatment. Brains were harvested and flash frozen in isopentane for molecular analysis.

### Molecular Analysis

#### Quantitative real-time PCR

Brains were sectioned into 1 mm thick coronal slices, and brain punches were collected from the PFc, NAc, and dorsal striatum. For cohort 1, punches were obtained from the bulk NAc. For cohort 2, NAc core (NAcC) and shell (NAcS) subregions were collected from one hemisphere. RNA was isolated using commercially available kits (Zymo Research, R1014) and cDNA was synthesized using 50 nanograms of RNA with qScript cDNA supermix (VWR Scientific, 10142-788). Probes from ThermoFisher Taqman for Oprm1, Oprk1, Pdyn, Penk, Grin2a, Grin2b and Gapdh (used for housekeeping) were utilized. Genes were run in a Roche LightCycler II. Samples were run in triplicate. Bulk NAc from the remaining cohort 2 hemispheres was grossly collected for protein- and signaling-level assays.

#### Membrane preparation

Membranes were prepared from NAc of individual saline and 10 mg/kg ketamine treated rats sacrificed 7 days later as described previously [13]. Briefly, tissue was homogenized with 25 volumes (1 g wet weight/25 ml) of ice-cold 20 mM Tris-Cl buffer (see Supplemental Material) and centrifuged at 27,000*g* for 15 min at 4°C. The pellet was resuspended in 25 ml of the same buffer, and the centrifugation step repeated. The final membrane pellet was resuspended in 40 volumes (of original wet weight) of 2 mM Tris-Cl buffer (see Supplemental Material). The protein content was determined using the Pierce BCA Protein Assay Reagent. Membrane preparations were used for ELISA and [^35^S]GTP γS binding assays.

#### ELISA

Levels of MOR protein were measured using ELISA with anti-MOR monoclonal antibody (5G8, Devi Laboratory [19]) as previously described [18]. Briefly, membranes (2 μg/100 μl 1x PBS) were coated on each well of a high-binding polystyrene 96-well plate. After the plate was dry, non-specific sites were blocked with 3% BSA in PBS for 1 h. Wells were incubated overnight at 4 ^°^C with the 5G8 antibody (1:500). Antibody solution was subsequently removed, and wells were washed three times (5 min/wash) with PBS containing 1% BSA followed by incubation with anti-mouse IgG coupled to horseradish peroxidase (cat. no. PI-2000, Vector Labs) for 90 min at RT. Antibody solution was removed, wells were washed three times (5 min/wash) with PBS containing 1% BSA and 200 *μ*l of the enzyme substrate, o-phenylenediamine (see Supplemental Material) was added to each well. The reaction was terminated within 3 – 5 min by the addition of 100 *μ*l 3 N HCl and absorbance measured at 490 nm with a Bio-Rad plate reader. More details in supplemental methods.

#### [^35^S] GTPγS binding

GTPγS binding assays were carried out as previously described [20]. Membranes (20 μg protein) were incubated for 1 hour at 30 °C in the absence (0 = 0.1% DMSO in assay buffer) or presence of DAMGO (10 *μ*M) in assay buffer (see Supplemental Material) containing freshly prepared 30 *μ*M GDP (for basal signal), and 0.1 nM [^35^S]GTP*γ*S. Nonspecific binding was determined in the presence of 10 *μ*M cold GTP*γ*S. Basal values represent values obtained in the presence of GDP and in the absence of ligand. At the end of the incubation period, samples were filtered using a Brandel filtration system. Filters were washed 3 times with 3 mL of ice-cold 50 mM Tris-Cl buffer, pH 7.4, and bound radioactivity was measured using a scintillation counter (MicroBeta TriLux; PerkinElmer).

#### Radioimmunoassay

Radioimmunoassays were carried out as described previously [22,23]. Briefly, NAc samples from cohort 3 animals were homogenized, acid-extracted, and dried overnight. Samples were subsequently resuspended in 500 μl of 50 mM sodium phosphate buffer, pH 7.6. To determine total Leu-enkephalin levels, a subset of samples was proteolytically cleaved with trypsin followed by carboxypeptidase B treatment to liberate Leu-enkephalin from larger precursor peptides as described [23]. For this, an aliquot (60 μL) of each fraction was treated with 5 μg/mL tosylphenylalanylchloromethyl ketone-treated trypsin (Sigma) for 16 h followed by treatment with 5 ng/mL carboxypeptidase B (CPB, Sigma) for 120 min, and the reaction was terminated by boiling for 20 min. The radioimmunoassay was carried out overnight at 4°C in a final volume of 300 μl in assay buffer A (see Supplemental Material) using samples and Leu-enk standards, rabbit anti-Leu-enkephalin IgG antibody (Phoenix Pharmaceuticals, H-024-21) and [^125^I]-Leu-enkephalin (Phoenix Pharmaceuticals, T-024-21). Bound radioactivity was separated from free by rapid filtration using 0.45 uM nitrocellulose filters [21] that were then washed 3 times with ice-cold 50 mM phosphate buffer pH 7.6. Levels of immunoreactive Leu-enkephalin were calculated by subtracting nonspecific binding and extrapolating from standard curve values using Prism software (GraphPad, San Diego, CA). More details in supplemental methods.

### Heroin Self-Administration Cohort

#### Animals and Surgical Procedures

A total of 26 adult male Long-Evans rats (8 weeks old) were included in the final study. Rats were anesthetized using isoflurane (5% induction, 3-4% maintenance) and implanted with jugular vein catheters as previously described [25].

#### Intravenous Heroin Self-Administration

Rats were trained to self-administer 30 μg/kg/infusion heroin under a fixed-ratio 1 schedule in operant chambers containing active and inactive levers [26]. Active lever presses resulted in heroin delivery paired with environmental cues, while inactive lever presses had no programmed consequences. Acquisition was defined as at least 2× more active than inactive lever presses. Rats then underwent 14 experimental sessions during which they self-administered heroin for 3 hours.

#### Forced Abstinence and Ketamine Interventions

After the self-administration portion, rats encountered 13 days of forced abstinence during which they remained in their home cages [26]. Rats received 3 intravenous infusions of 2.35 mg/kg ketamine dissolved in 0.9% saline (Ketasthesia, racemic, Henry Schein) delivered intermittently over 40 minutes on days 9, 11, and 13 of forced abstinence in modified operant chambers with white wallboards to create a distinct context. Rats also received a 10 mg/kg ip dose of ketamine 15 days after the final heroin self-administration session.

#### Heroin-Seeking Activities

Rats underwent 3 heroin-seeking sessions [26]. The first two (days 14 and 18 post-self-administration) were 1-hour cue-induced sessions during which all parameters remained the same, but heroin was unavailable. The final session was heroin-primed: rats received 0.25 mg/kg ip heroin 30 minutes prior to the task, 7 days after the 10 mg/kg ip ketamine injection. Rats were sacrificed 90 minutes after this final session and brains were flash frozen in isopentane for molecular analyses.

### Statistical Analyses

Statistical analyses were performed using GraphPad Prism (version 10.4.1). For qPCR data, fold change was calculated using the 2^-ΔΔCt^ method with GAPDH as the reference gene. Fold changes were analyzed using two-way or mixed-models ANOVA to evaluate interactions between timepoint and drug treatment, followed by Tukey’s post-hoc analysis. ELISA data were analyzed using t-tests. For GTPγS assays, counts per minute (cpm) were analyzed using two-way ANOVA to evaluate interactions between ketamine treatment and DAMGO, followed by Tukey’s post-hoc analysis. GTPγS data were then normalized to basal conditions and analyzed using a t-test. For RIA, one-way ANOVA followed by Dunnett’s post-hoc analysis was utilized. For the self-administration cohort, repeated-measures ANOVA confirmed acquisition. Additional ANOVA tests analyzed heroin-seeking activities with Sidak’s post-hoc tests. Pearson correlation matrices assessed interactions between behavioral and qPCR data.

## Results

### Acute ketamine alters transcription of genes associated with the endogenous opioid system in the NAc in a time- and dose-dependent manner

To investigate ketamine’s molecular mechanisms, a cohort of animals underwent an acute administration of two doses of ketamine, followed by assessment of endogenous opioid gene transcription using qPCR (Fig 1A). In the NAc, main effects of Treatment were observed for Oprm1 (F2,9=6.41, p=0.019) (Fig 1B), Oprk1 (F2,9=42.34, p<0.001) (Fig 1C), Pdyn (F2,9=12.92, p=0.002) (Fig 1D), and Penk (F2,9=20.56, p<0.001) (Fig 1E). Main effects of Time were observed for Oprk1 (F1.68,14.26=27.13, p<0.001), Pdyn (F1.695,14.40=14.73, p<0.001), and Penk (F1.53,13.03=13.10, p=0.001). Significant Treatment × Time interactions were observed for all four genes (Oprm1: F4,17=3.51, p=0.029; Oprk1: F4,17=10.19, p<0.001; Pdyn: F4,17=5.29, p=0.006; Penk: F4,17=4.49, p=0.012). Post hoc analysis revealed that both 3 mg/kg and 10 mg/kg ketamine significantly increased transcription of all four genes at the 3-day timepoint (p<0.02) (Fig 1B-E). These effects were sustained at the 7-day timepoint only in the 10 mg/kg group (Oprm1: p=0.003; Oprk1: p=0.004; Pdyn: p=0.001; Penk: p=0.020) (Fig 1B-E), demonstrating synergistic time- and dose-dependent effects of ketamine across all four endogenous opioid system related genes assessed in the NAc. No significant changes were observed in the PFc or dorsal striatum (Supp Fig 1A and 1B).

**Figure 1.**
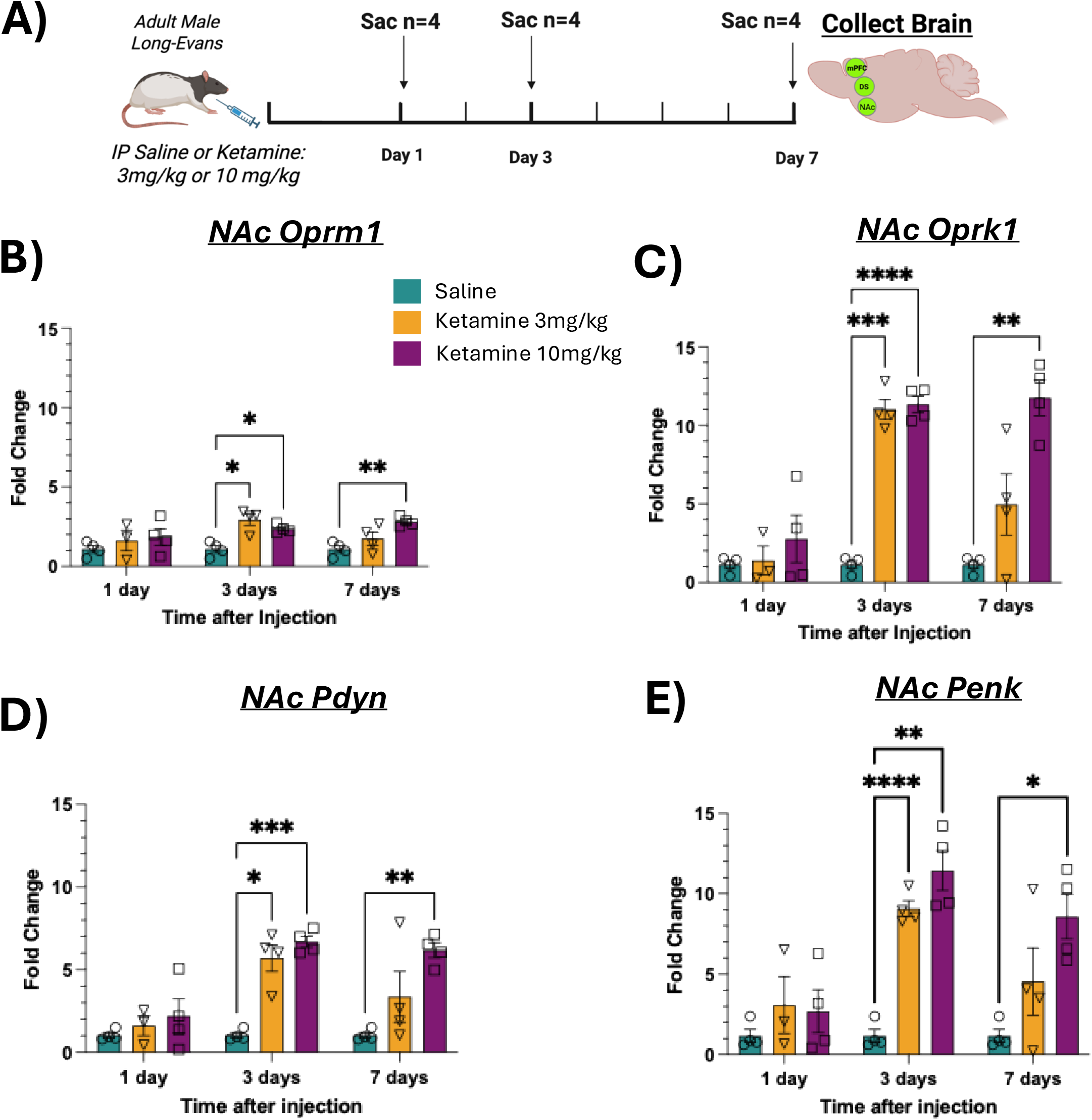
Acute ketamine produces dose- and time-dependent upregulation of endogenous opioid system genes in the nucleus accumbens. (A) Experimental design. Adult male Long-Evans rats received saline, 3 mg/kg, or 10 mg/kg ketamine intraperitoneally and were sacrificed at 1, 3, and 7 days post-injection (n=4/group/timepoint). Whole nucleus accumbens (NAc), medial prefrontal cortex (PFc), and dorsal striatum (DS) were collected for qRT-PCR. (B) NAc *Oprm1*, (C) NAc *Oprk1*, (D) NAc *Pdyn*, and (E) NAc *Penk* mRNA were significantly upregulated at the 3-day timepoint in both dose groups, with effects sustained at the 7-day timepoint only in the 10 mg/kg group. *p<0.05, **p<0.01, ***p<0.001, ****p<0.0001. Data are mean ± SEM.

### 10mg/kg ketamine alters transcription of genes associated with the endogenous opioid system in the NAc shell but not core

Next, as the NAc consists of functionally distinct subregions playing differing roles in addiction, we investigated NAc core (NAcC) and NAc shell (NAcS) subregional transcriptional differences in the second acute ketamine cohort (Fig 2A). In the NAcC, no main effects on the transcription of Oprm1, Oprk1, Pdyn or Penk were observed (Fig 2B-E). However, post-hoc analysis revealed Oprm1 transcription was increased at the 1-day timepoint (p=0.011) (Fig 2B). In the NAcS, main effects of Treatment were observed for Oprm1 (F1,13=6.99, p=0.020) (Fig 2F), Oprk1 (F1,13=4.89, p=0.046) (Fig 2G), Pdyn (F1,13=4.81, p=0.047) (Fig 2H), and Penk (F1,13=6.84, p=0.021) (Fig 2I). Main effects of Time were observed for Oprk1 (F2,24=4.96, p=0.016) and Penk (F1.78,22.20=4.05, p=0.036). Significant Time × Treatment interactions were observed for Oprk1 (F2,24=4.96, p=0.016) and Penk (F2,25=4.05, p=0.030). Post hoc analysis revealed transcriptional increases of Oprm1 at the 1-day (p=0.027) and 7-day (p=0.033) timepoints (Fig 2F), Oprk1 and Pdyn (p=0.003, p=0.017) (Fig 2G, Fig 2H) at the 7-day timepoint, and Penk at the 3-day (p=0.022) and 7-day (p=0.007) timepoints (Fig 2I), highlighting NAcS as the key driver of ketamine-induced endogenous opioid system changes.

**Figure 2.**
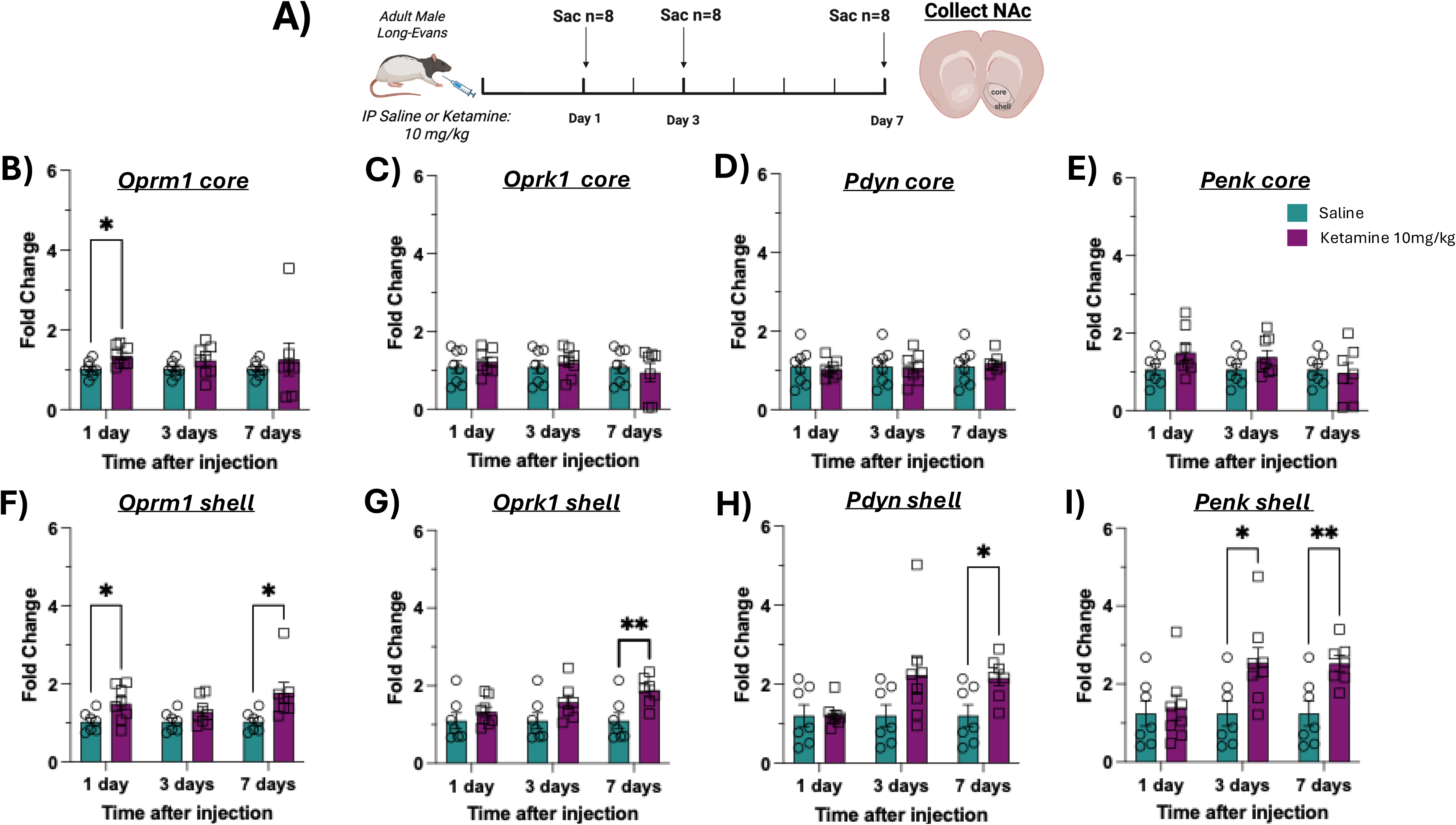
Ketamine-induced opioid gene upregulation is restricted to the nucleus accumbens shell. (A) Experimental design. Rats received saline or 10 mg/kg ketamine ip and were sacrificed at 1, 3, and 7 days post-injection (n=8/group/timepoint). NAc core (NAcC) and shell (NAcS) were microdissected for qRT-PCR. (B–E) In the NAcC, no significant main effects of ketamine were observed for (B) *Oprm1*, (C) *Oprk1*, (D) *Pdyn*, or (E) *Penk*, though *Oprm1* was transiently increased at the 1-day timepoint. (F–I) In the NAcS, (F) *Oprm1*, (G) *Oprk1*, (H) *Pdyn*, and (I) *Penk* were all significantly elevated at the 7-day timepoint. *p<0.05, **p<0.01. Data are mean ± SEM.

### Acute ketamine alters transcription of genes associated with the glutamatergic systems in the PFc and NAc subregions

Next, to probe for potential ketamine-induced plasticity mechanisms, we investigated transcriptional changes to Grin2a and Grin2b, which encode NMDA receptor subunits implicated in plasticity. No significant changes were observed in brain regions assessed in the first acute cohort (Supp Fig 2A and B). However, in the second cohort (Fig 3A) a main effect of Treatment was observed for Grin2b (F1,14=9.31, p=0.009) in the PFc (Fig 3B), with decreased transcription at the 1-day (p=0.031) and 3-day (p=0.004) time points. No changes to Grin2a were observed (Fig 3C). In the NAcC, main effects of Treatment were observed for Grin2b (F1,14=4.65, p=0.049) (Fig 3D) and Grin2a (F1,14=5.60, p=0.033) (Fig 3E). Grin2b transcription increased at the 3-day timepoint (p=0.013) whereas Grin2a increased at the 7-day timepoint (p=0.043). Interestingly, in the NAcS, no changes were observed in Grin2b (Fig 3F). A main effect of Time (F2,25=4.55, p=0.021) and a Time × Treatment interaction (F2,25=4.55, p=0.021) were detected for Grin2a (Fig 3G), with increases at the 3-day (p=0.045) and 7-day (p=0.034) time points. The absence of glutamatergic changes in the first cohort likely reflects its smaller sample size and use of bulk NAc tissue, whereas the larger sample size and subregional (core versus shell) dissection in the second cohort afforded greater sensitivity to detect these relatively modest, subregion-specific transcriptional effects.

**Figure 3.**
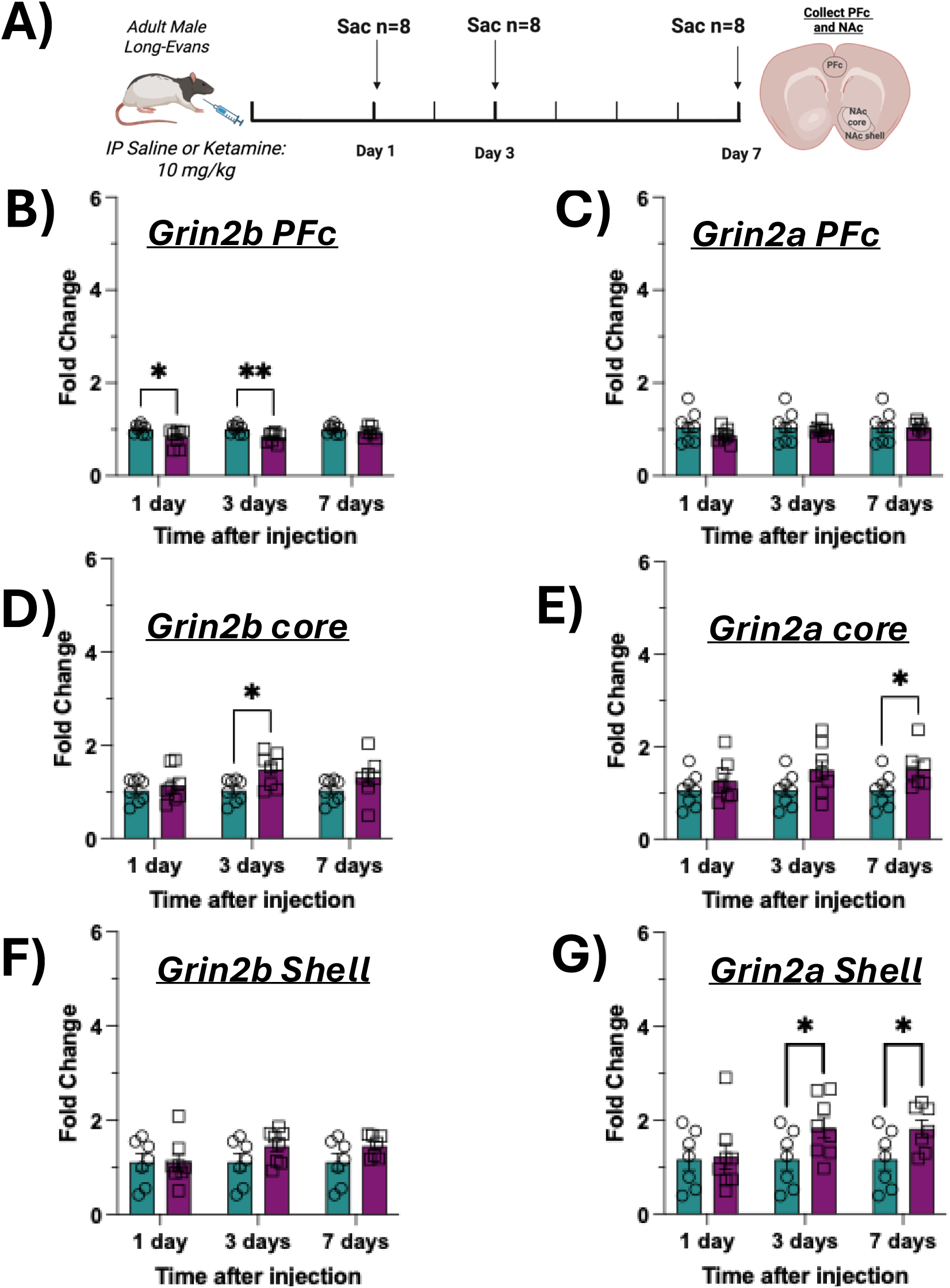
Ketamine produces region- and subunit-selective changes in NMDA receptor gene expression. (A) Experimental design. Rats received saline or 10 mg/kg ketamine ip and were sacrificed at 1, 3, and 7 days post-injection (n=8/group/timepoint). PFc, NAcC, and NAcS were dissected for qRT-PCR of *Grin2a* and *Grin2b*. (B–C) In the PFc, (B) *Grin2b* was downregulated at the 1- and 3-day timepoints (*p<0.05, **p<0.01); (C) *Grin2a* was unchanged. (D–E) In the NAcC, (D) *Grin2b* was increased at the 3-day timepoint and (E) *Grin2a* at the 7-day timepoint (*p<0.05). (F–G) In the NAcS, (F) *Grin2b* was unchanged; (G) *Grin2a* was elevated at the 3- and 7-day timepoints (*p<0.05). Data are mean ± SEM.

### 10mg/kg ketamine alters endogenous opioid system at the protein level in the NAc 7 days after acute administration

As the 10 mg/kg dose of ketamine and 7-day time point elicited the most robust transcriptional alterations to the NAc endogenous opioid system, we next investigated whether these changes translated to the protein level (Fig 4A). Using immunoassays to quantify MOR protein levels (Fig 4B), we observed increased MOR abundance in the NAc of ketamine-treated animals from cohort 1 (p=0.009) (Supp Fig 3A). Importantly, this increase in MOR abundance was also observed in a second acute ketamine cohort, consisting of a larger number of animals (p<0.001) (Fig 4C). Next, to assess if the observed MOR upregulation was accompanied by changes in receptor signaling, we used a G protein activity assay to measure DAMGO-mediated increases in GTPγS binding in the second acute ketamine cohort (Fig 4D). Raw counts per minute (cpm) revealed a main effect of ketamine (F1,28=7.983, p=0.008) and DAMGO (F1,28=135.2 p<0.0001) treatment on increases in GTPγS binding (Supp Fig 3B). Data normalized to basal conditions showed a main effect of ketamine (F1,28=50.01 p<0.0001), DAMGO (F1,28=1033 p<0.001) and interaction (F1,28=50.01 p<0.001) on GTPγS binding (Fig 4E), indicating the 10 mg/kg dose of ketamine enhances GTPγS binding following MOR activation 7 days post treatment. Next, we used radioimmunoassays to directly quantify levels of endogenous opioid peptides, namely Leu-Enkephalin (Leu-Enk) 7 days post treatment with 10 mg/kg ketamine. The results revealed a main effect of ketamine treatment on Leu-enk levels in the NAc (p<0.001) (Fig 4F-G), with significant increases in both free (p<0.001) and total Leu-enk (p<0.001); the total Leu-Enk was assessed following trypsin/CPB treatment - a treatment that is known to release Leu-enk from the larger Leu-enkephalin-containing peptides such as proDyn, Neo-endorphins as well as proEnk-derived Peptide E [24] Together, these results show that ketamine treatment increases MOR levels and signaling and Leu-Enk levels (free & total) in the NAc.

**Figure 4.**
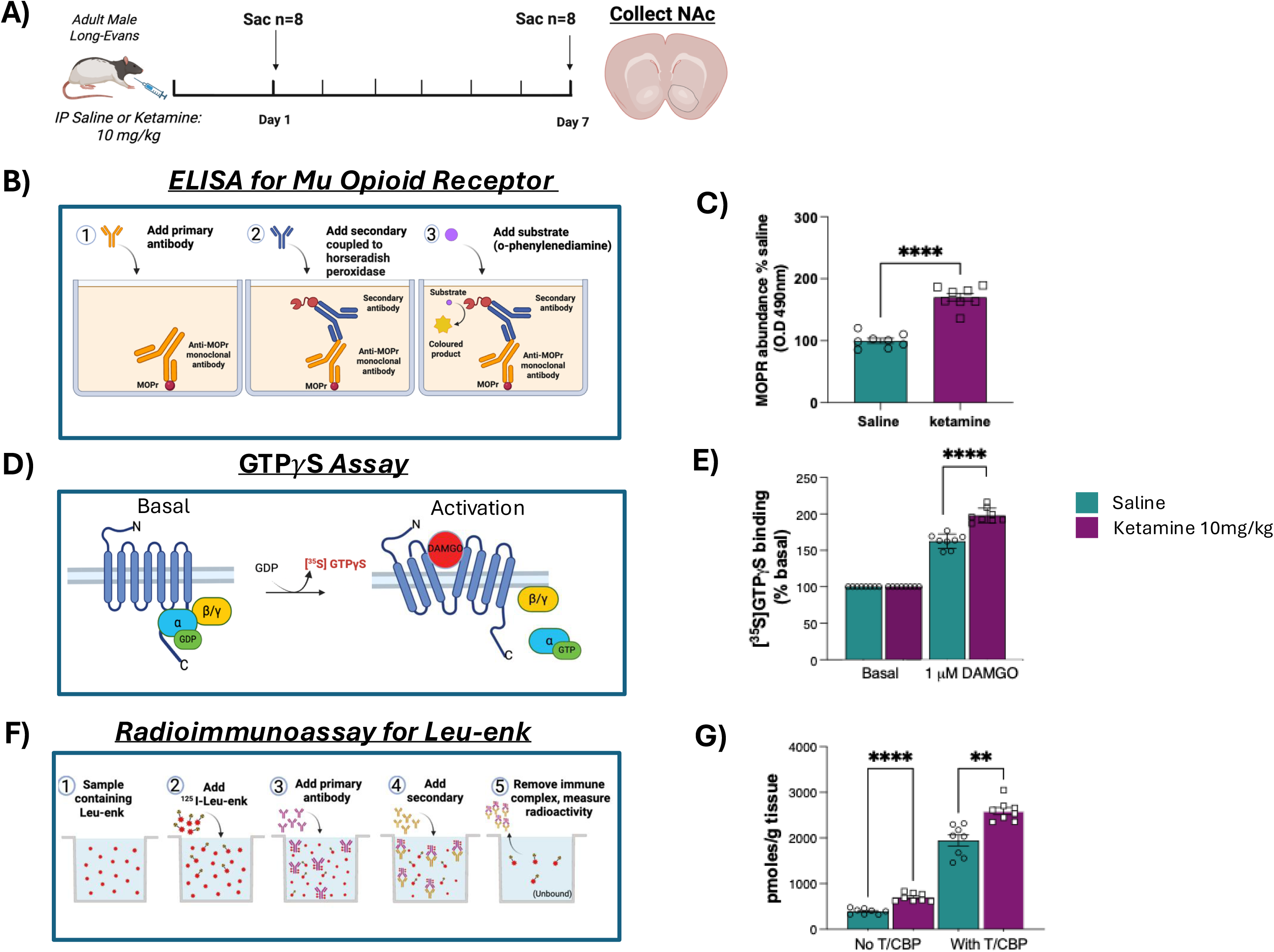
Ketamine increases mu-opioid receptor protein, G-protein coupling, and Leu-enkephalin peptide levels in the NAc. (A) Experimental design. Rats received saline or 10 mg/kg ketamine ip and were sacrificed at the 7-day timepoint (n=8/group). Whole NAc was processed for ELISA, [^35^S]GTPγS binding, and radioimmunoassay (RIA). (B) Schematic of ELISA approach. (C) ELISA revealed significantly elevated MOR protein levels in the NAc of ketamine-treated animals (****p<0.0001). (D) Schematic of GTPγS binding assay. (E) [^35^S]GTPγS binding demonstrated enhanced DAMGO-stimulated receptor-G-protein coupling in the NAc following ketamine treatment (****p<0.0001). (F) Schematic of RIA approach. (G) RIA demonstrated significantly increased free and total Leu-enkephalin peptide levels in ketamine-treated animals (****p<0.0001, **p<0.01). Data are mean ± SEM.

### Ketamine interventions decrease heroin-primed seeking activity following a forced abstinence period

Finally, to assess ketamine’s potential to decrease relapse in OUD, a third cohort of animals underwent a heroin self-administration paradigm (Fig 5A). Animals received ketamine during abstinence followed by three sessions assessing heroin-seeking activity. No significant effects were observed during the first cue-induced session (Fig 5B). Following a 10 mg/kg ip ketamine dose, the second cue-induced session revealed a main effect of Treatment (F1,24=6.06, p=0.021), with ketamine-treated animals pressing the inactive lever fewer times (p=0.007) (Supp Fig 4A). For the heroin-primed session 7 days post-ketamine, Sidak’s correction revealed that ketamine-treated animals pressed the active lever significantly fewer times than saline-treated animals (p=0.036) (Fig 5C). No significant transcriptional alterations were observed in the NAcC (Supp Fig 5A-E), NAcS (Supp Fig 6A-F), or PFc (Supp Fig 7A-F) following behavioral testing. Pearson correlations revealed no significant associations in NAcC of saline or ketamine-treated animals (Supp Fig 8A-B) or NAcS of saline-treated animals (Supp Fig 8C). However, in the NAcS of ketamine-treated animals, active lever presses correlated with Pdyn (r=0.73, p=0.012), and infusions correlated with Pdyn (r=0.81, p<0.001) and Penk (r=0.69, p=0.01) (Supp Fig 8D).

**Figure 5.**
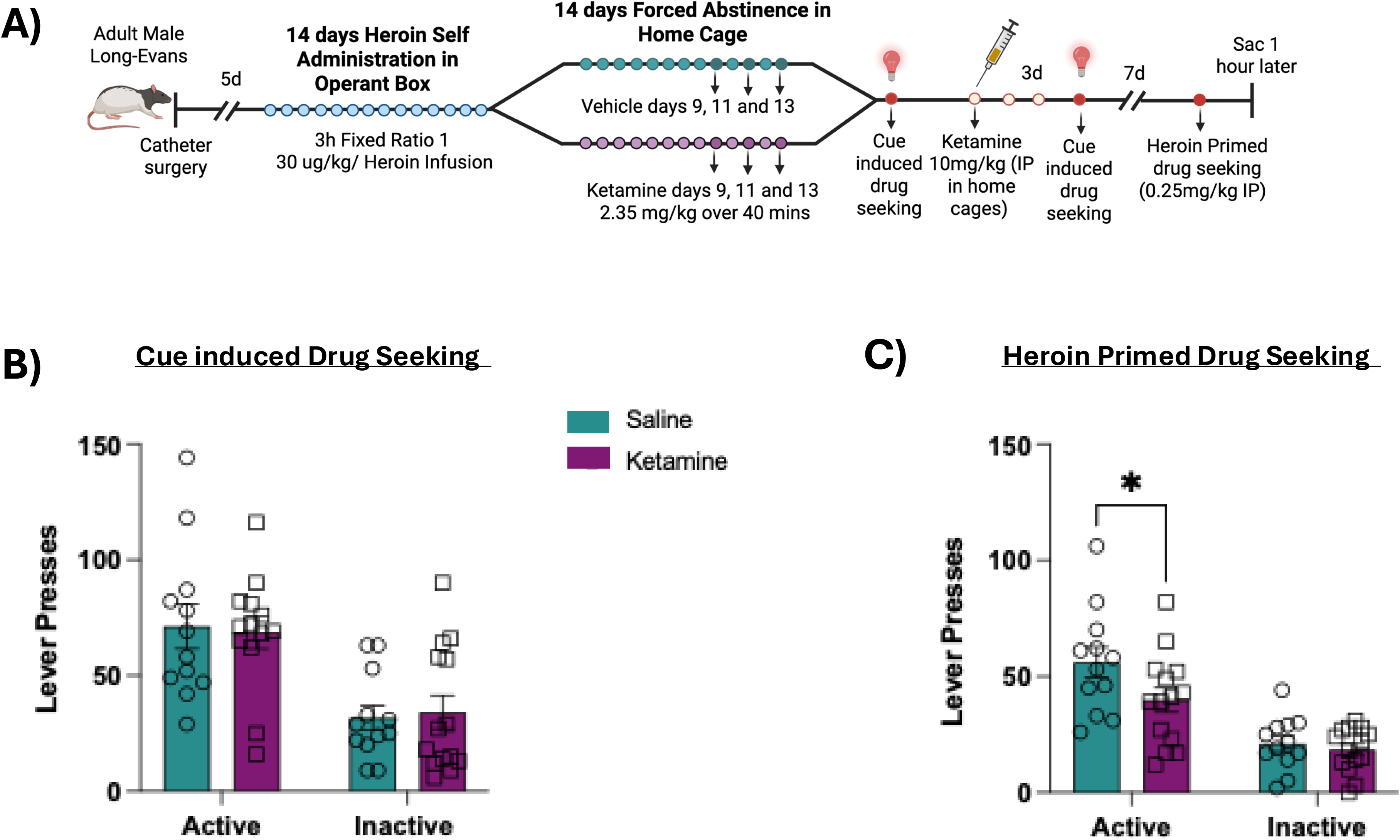
Sub-anesthetic ketamine reduces heroin-primed but not cue-induced drug seeking. (A) Experimental timeline. Rats underwent heroin self-administration (30 μg/kg/infusion, FR1, 14 days) followed by forced abstinence with vehicle or ketamine treatment (2.35 mg/kg iv, days 9, 11, and 13) and a 10 mg/kg ip ketamine dose on day 15. Cue-induced and heroin-primed reinstatement were assessed sequentially. (B) Ketamine did not significantly alter active or inactive lever pressing during the first cue-induced drug-seeking session. (C) Ketamine significantly reduced active lever pressing during heroin-primed reinstatement 7 days post-ip injection (*p<0.05). Data are mean ± SEM.

## Discussion

The present study demonstrates that acute ketamine remodels endogenous opioid circuitry in the NAc at the transcriptional, protein, and MOR signaling levels, and reorganizes NMDAR subunits implicated in plasticity. Behaviorally, ketamine decreased heroin-primed seeking when administered in a therapeutically relevant timeframe.

Transcription of genes encoding MOR, KOR, prodynorphin, and proenkephalin increased in the NAc 7 days after a single ketamine administration, suggesting sustained opioid system reorganization in addiction-related regions. These alterations were accompanied by increased MOR protein and enhanced MOR-mediated G-protein activation, together with elevated proDyn and proEnk mRNA and Leu-enkephalin peptide (total and free), indicating enhanced precursor synthesis and processing. Although similar long-lasting opioid system adaptations have not been reported, 10 mg/kg ip ketamine has previously been shown to increase Oprm1, POMC, and β-endorphin levels [9]. Because ketamine orthosterically binds MOR [8] and acts as a partial agonist, the observed multi-faceted effects may initially arise through direct receptor engagement. However, ketamine is rapidly metabolized to 6-hydroxynorketamine, a major bioactive metabolite detected in rodent [11] and human [12] plasma, suggesting its sustained effects are driven by metabolites. Because MOR binding is limited to ketamine but not its major metabolites [28], the sustained molecular adaptations are likely mediated through orthosteric binding-independent mechanisms. Interestingly, ketamine and 6-hydroxynorketamine also act as opioid receptor PAMs, enhancing endogenous opioid peptide-driven G-protein signaling [10, 29]. We propose that this acute allosteric engagement is followed by a slower phase of transcriptional regulation culminating in the protein- and signaling-level changes observed here.

Notably, proenkephalin is among the most highly upregulated genes following chronic antidepressant treatment; however, this induction typically requires 14 or more days of daily fluoxetine dosing to elevate enkephalin peptide and Penk levels [29]. That a single dose of ketamine is sufficient to upregulate Penk and Leu-enkephalin lasting up to 7 days distinguishes it from conventional antidepressants and may reflect its additional PAM activity at MOR. These long-lasting adaptations are particularly relevant in OUD, where chronic opioid exposure diminishes receptor responsiveness through a reduction in coupling to G proteins, drives receptor internalization [30], and reduces Penk and Pdyn mRNA expression [31]. By upregulating MOR expression, opioid peptide levels, and MOR-mediated G-protein activation, ketamine may restore disrupted opioid signaling in OUD and reduce relapse.

The transcriptional increases observed were specific to the NAcS but not NAcC. The NAcS efferents project to the lateral hypothalamus and amygdala [32] — regions mediating stimulus-induced drug seeking and affective-driven behaviors [33]. The NAcS is also more sensitive to ventral tegmental area dopaminergic input, rendering it key for drug-induced plasticity [34], and the PFc-NAcS circuit is necessary for context-induced heroin seeking [35].

Ketamine significantly reduced heroin-primed seeking activity following forced abstinence, supporting its potential to reduce relapse following acute opioid re-exposure. Although no other studies have utilized a heroin self-administration paradigm to investigate ketamine as an intervention for OUD, our findings are consistent with work showing NMDAR antagonism suppresses heroin-primed seeking after forced abstinence [27], and that ketamine attenuates morphine-conditioned place preference at similar doses [16,17].

Interestingly, endogenous opioid gene expression was not significantly altered in the self-administration cohort, suggesting differential effects when heroin is onboarded. Nonetheless, heroin-taking activity positively correlated with prodynorphin and proenkephalin expression in the NAcS of only ketamine-treated animals, suggesting ketamine enhances the functional relevance of the endogenous opioid system to heroin-seeking. Notably, the centrality of the opioid system to ketamine’s therapeutic actions was recently reinforced by work showing that ketamine’s rapid antidepressant effects are initiated by MORs enriched in somatostatin-expressing interneurons of the PFc, engaged through Gi/o signaling within minutes of administration [28]. Although that study localized MOR dependence to cortical interneurons, our findings extend ketamine-MOR engagement to the NAc and opioid relapse, suggesting MOR signaling is a shared feature of ketamine’s therapeutic actions.

Ketamine also produced region-specific alterations in transcription of Grin2a and Grin2b, the genes encoding NMDAR subunits GluN2A and GluN2B, respectively, suggesting synaptic remodeling within relapse-associated circuits. Activation of PFc-NAcS glutamatergic projections promotes heroin relapse [35], and GluN2b-mediated synaptic potentiation in this circuit is a key driver of this process. Accordingly, inhibition of GluN2b prior to heroin-primed seeking has been shown to block this potentiation, decreasing heroin-primed seeking activity [27]. We observed decreased Grin2b transcription in the PFc and increased Grin2b transcription in the NAcC at the 3-day timepoint, consistent with a role for this subunit in ketamine-induced plasticity. Interestingly, we also observed increased Grin2a expression in both the NAcC and NAcS. As the GluN2a/GluN2b ratio confers NMDARs with distinct pharmacological properties [36], our results suggest that ketamine alters NMDAR-subunit composition in this circuit, potentially normalizing glutamatergic signaling through a shift toward GluN2a-containing receptors. This reorganization may counteract the GluN2b-dependent potentiation that underlies heroin relapse.

Several limitations should be acknowledged. The sequential reinstatement design may have progressively reduced heroin-seeking independent of ketamine treatment, and future studies should evaluate heroin-primed reinstatement in isolation. Future studies should determine whether these mechanisms extend to other opioids such as fentanyl, examine KOR protein and signaling, and establish whether the NMDAR transcriptional changes translate to protein and signaling levels. Finally, evaluating these mechanisms in female animals is important, given known sex differences in ketamine pharmacokinetics [37]. We did not quantify proopiomelanocortin (Pomc); because Pomc is expressed at very low levels outside the hypothalamus and brainstem, its transcripts fell below the sensitivity of our qPCR assay in the regions examined. Future studies using more sensitive approaches will be needed to determine whether ketamine also modulates Pomc-derived β-endorphin signaling.

In conclusion, ketamine induces coordinated remodeling of endogenous opioid and glutamatergic signaling while reducing heroin-primed seeking following abstinence. By identifying transcriptional, protein-level, peptide, and functional adaptations within relapse-associated circuitry, these findings provide mechanistic insight into ketamine’s therapeutic efficacy against relapse in OUD.

## Data Availability Statement

The data that support the findings of this study are available from the corresponding author upon reasonable request.

## Acknowledgements

Special thanks to Maria Savoia, Daniel Garcia, Joseph Landry, James Callens, Samuel Cartwright, Alex Chisholm, Jacquie Ferland, Kion Winston, Katie Lynch, Alfonso Brea Guerrero, and Devin Hagarty for technical assistance and feedback.

## Author Contributions

**Adam M. Dawoud**: Writing – original draft, Data collection—behavioral experiments and qPCRs, Data curation, Conceptualization. **Achla Gupta:** Data collection— ELISAs, GTPγS assay and radioimmunoassays. **Ivone Gomes**: Data collection — radioimmunoassays, Writing – review & editing. **Dylan Baker:** Data curation, Writing – review & editing. **Julia Shapiro**: Data curation, Writing – review & editing. **Lakshmi Devi**: Supervision, Writing – review & editing, Funding acquisition and Conceptualization. **Yasmin L. Hurd**: Writing – review & editing, Supervision, Project administration, Funding acquisition, Data curation, Conceptualization. **Aya Osman**: Writing –original draft, review & editing, Data collection—ELISAs, GTPγS assay and radioimmunoassays, Data curation, Conceptualization, Supervision and Funding acquisition.

---

**Supplemental Figure 1.**
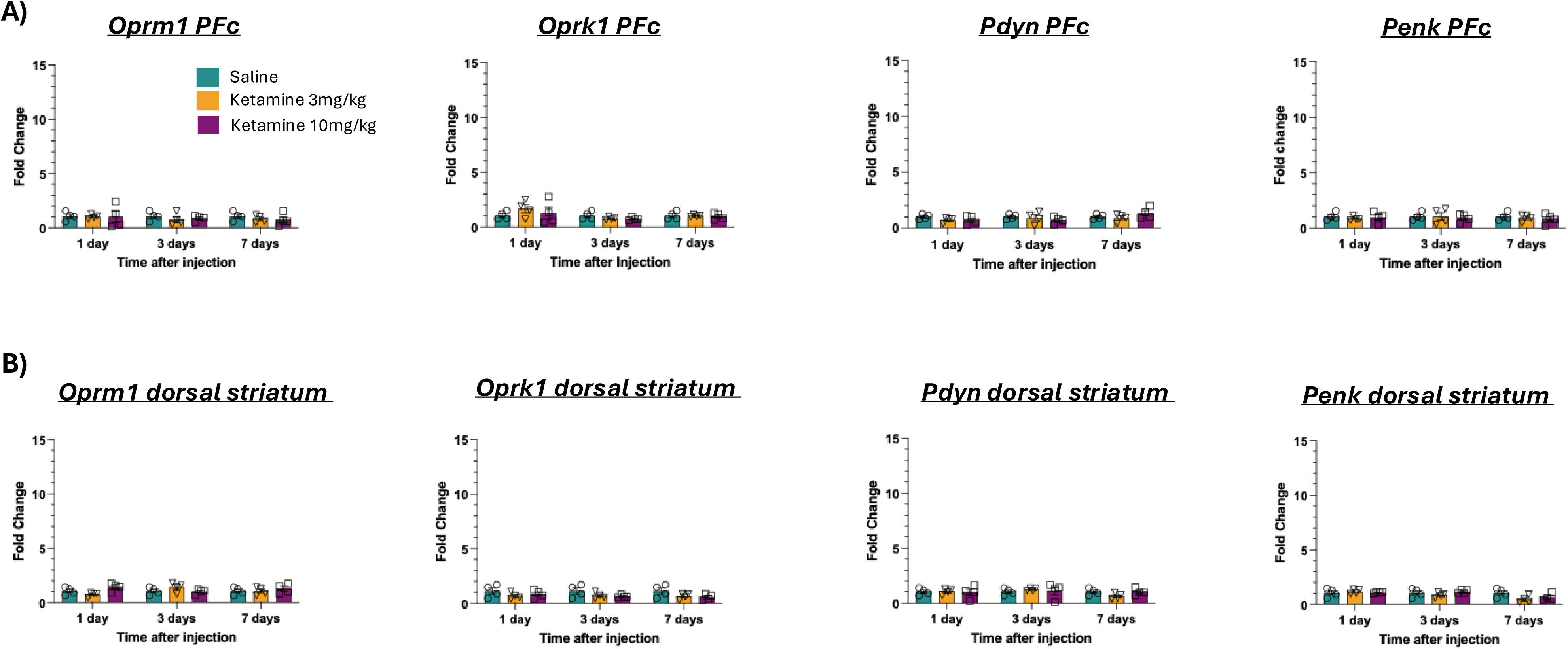

**Supplemental Figure 2.**
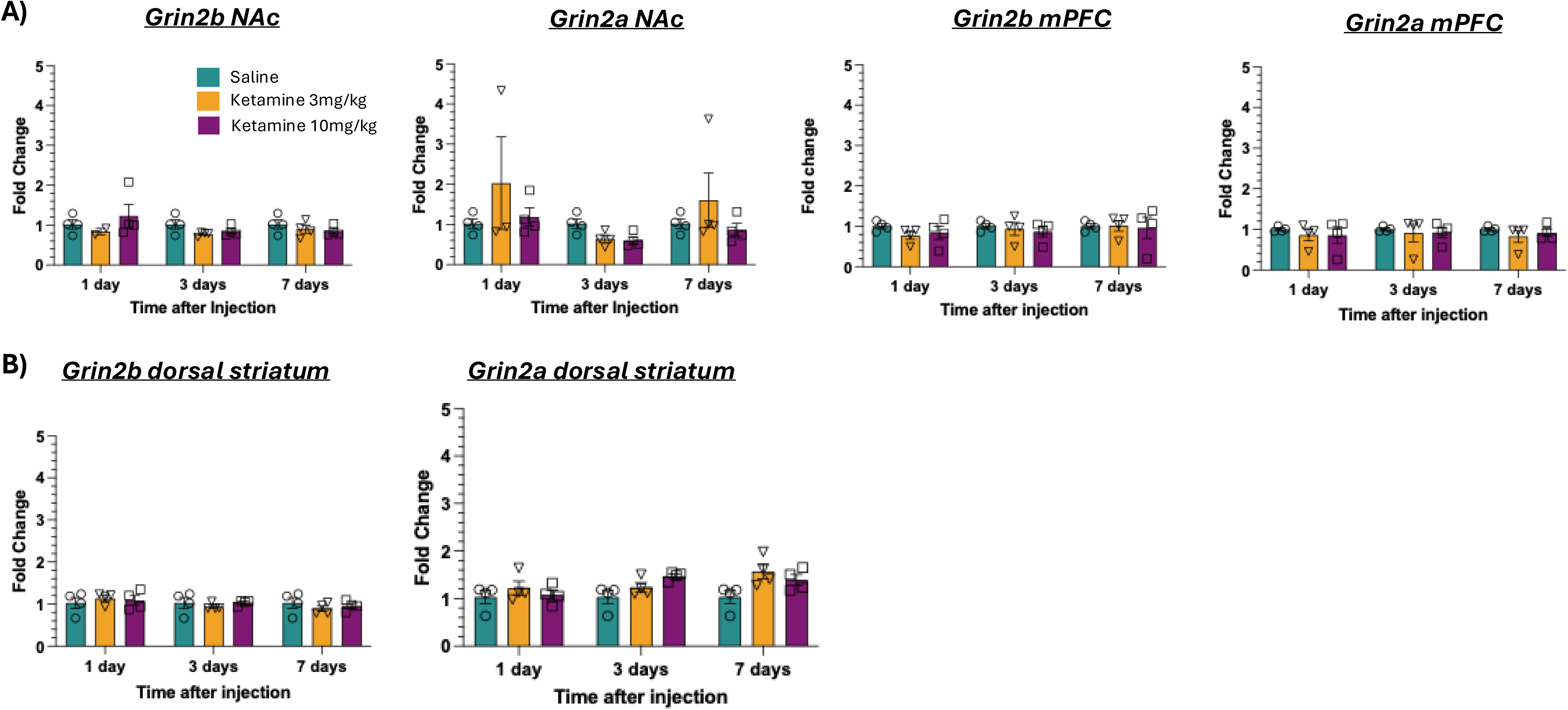

**Supplemental Figure 3.**
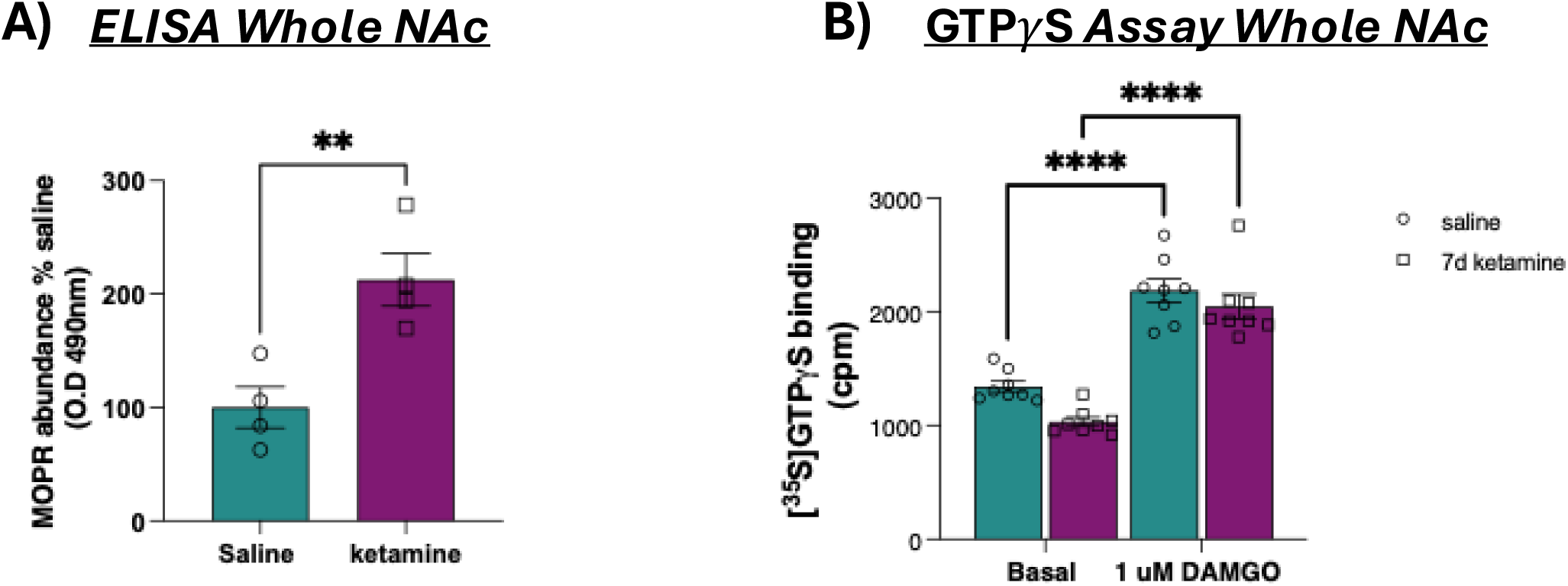

**Supplemental Figure 4.**
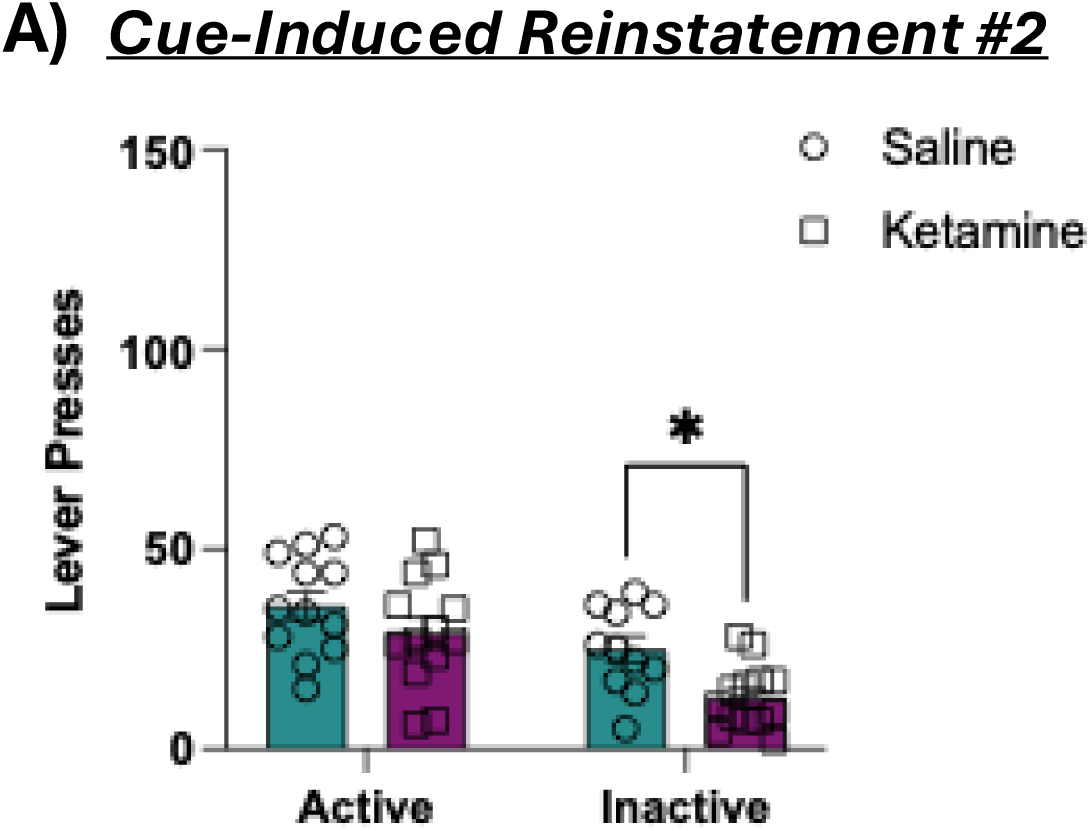

**Supplemental Figure 5.**
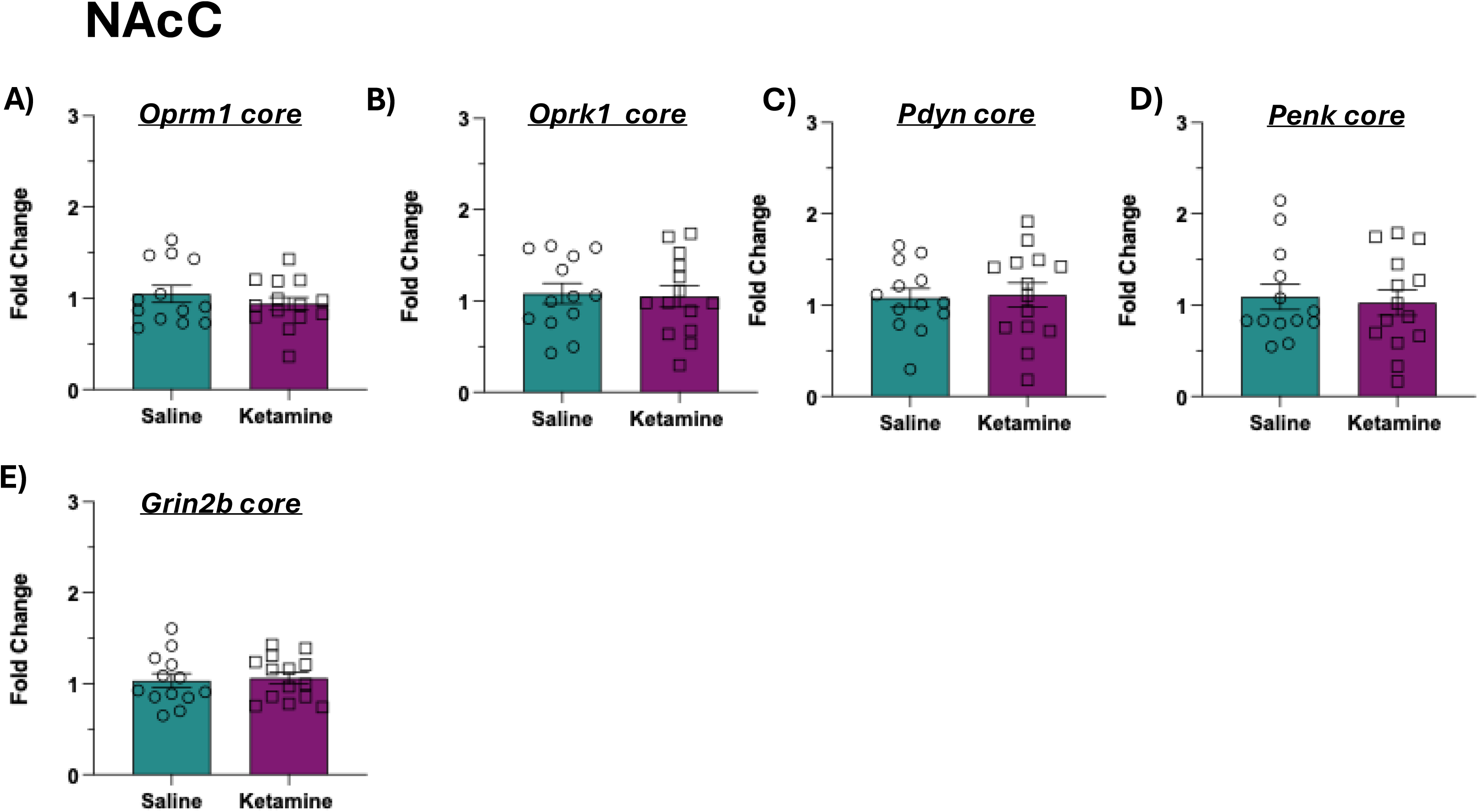

**Supplemental Figure 6.**
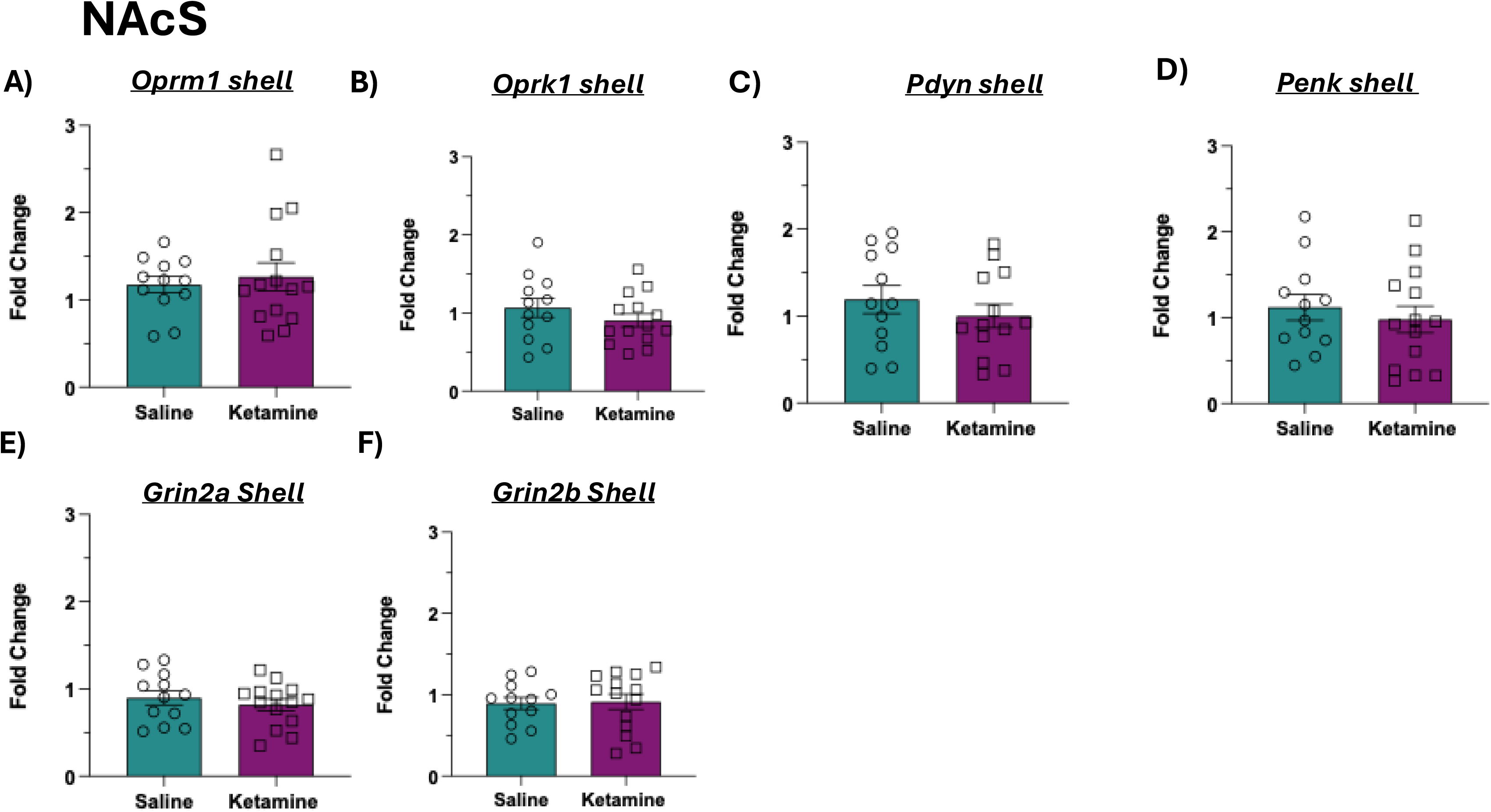

**Supplemental Figure 7.**
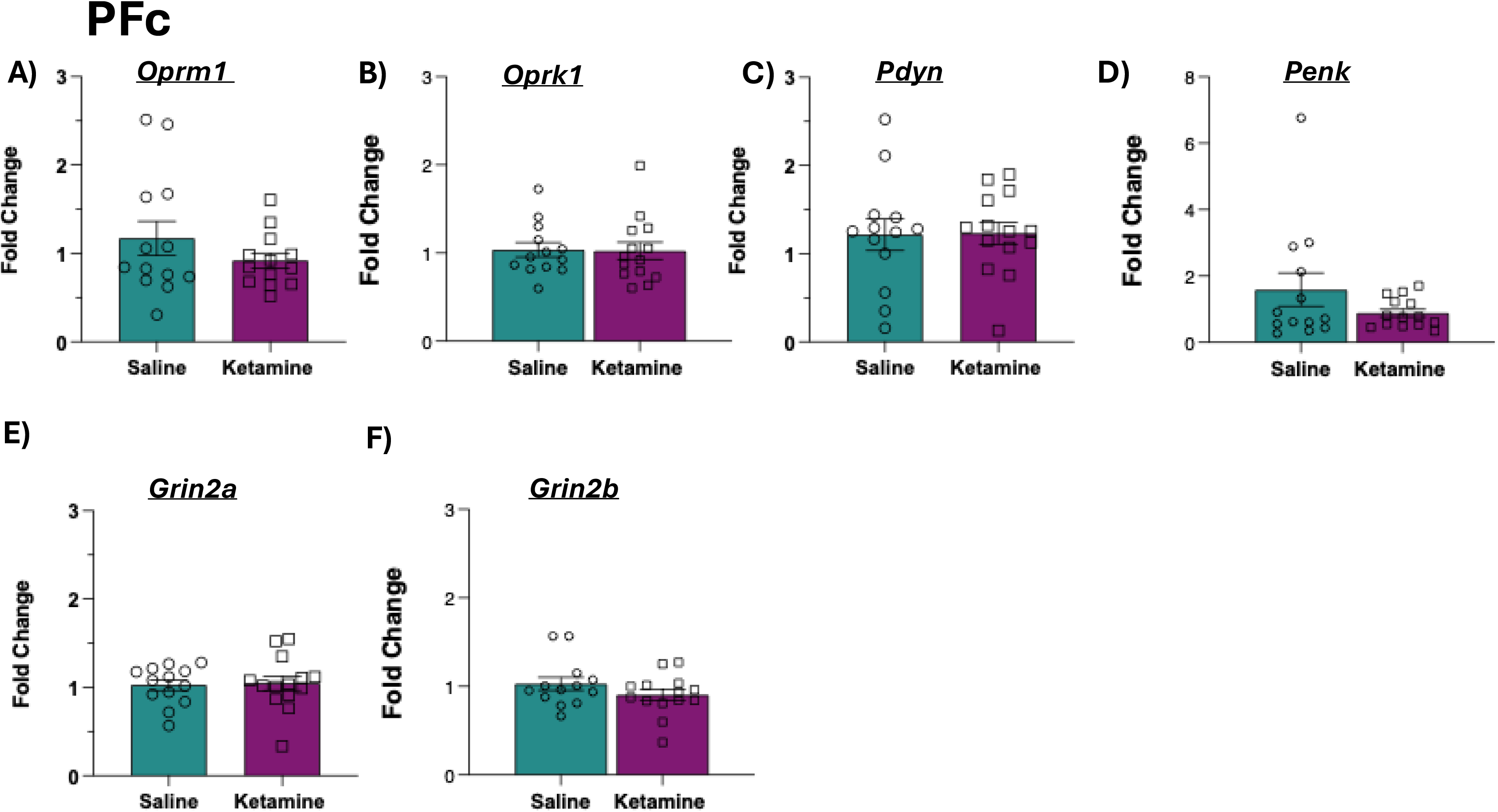

**Supplemental Figure 8.**
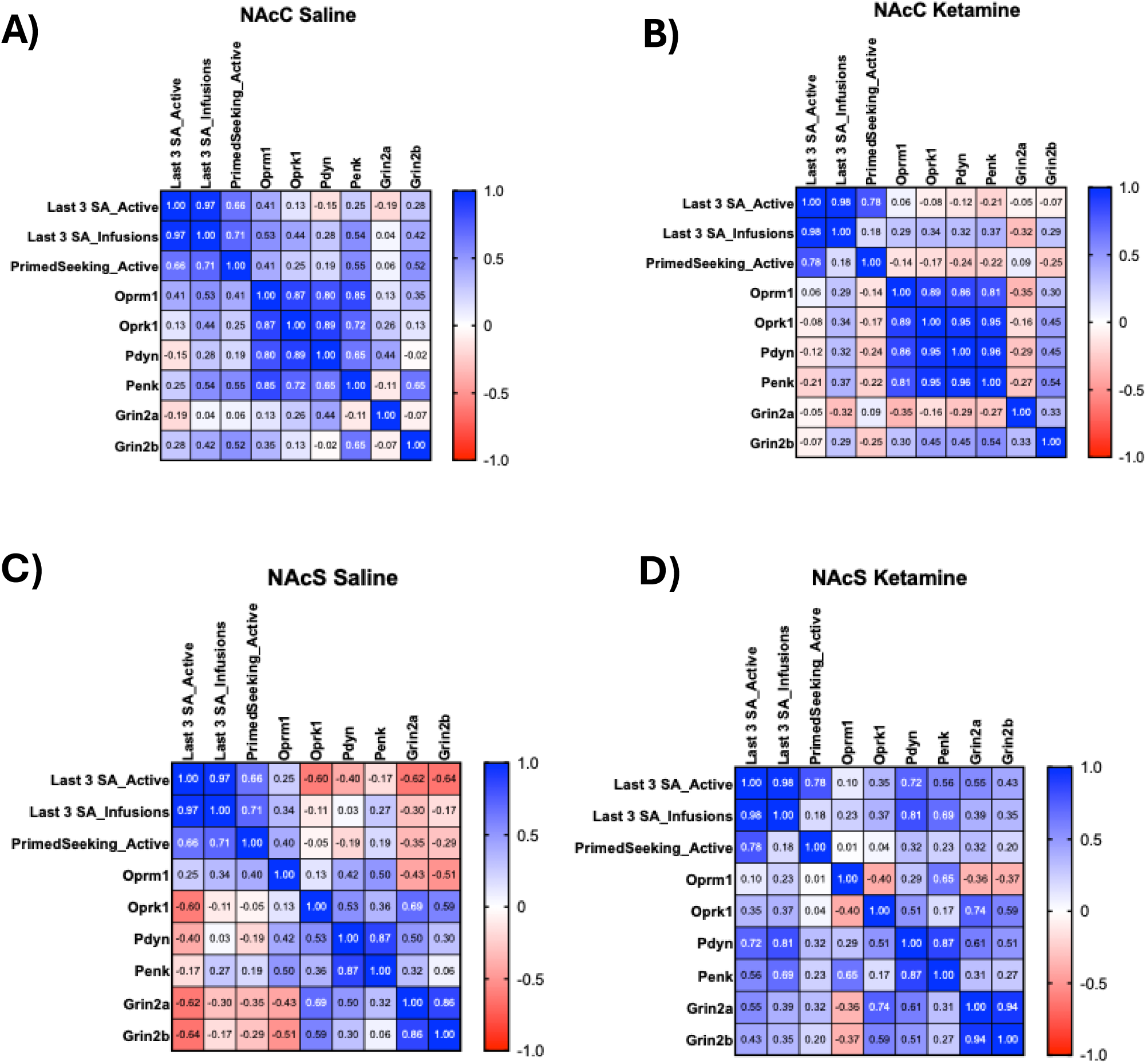

## References

1. Dydyk AM, Jain NK, Gupta M. Opioid Use Disorder. In: StatPearls. Treasure Island (FL): StatPearls Publishing; June 21, 2022.

2. Centers for Disease Control and Prevention (CDC). Understanding the epidemic. 2020. Retrieved from https://www.cdc.gov/drugoverdose/epidemic/index.html

3. Berman RM, Cappiello A, Anand A, Oren DA, Heninger GR, Charney DS, et al. Antidepressant effects of ketamine in depressed patients. Biol Psychiatry. 2000;47(4):351–354.

4. Aleksandrova LR, Phillips AG, Wang YT. Antidepressant effects of ketamine and the roles of AMPA glutamate receptors and other mechanisms beyond NMDA receptor antagonism. J Psychiatry Neurosci. 2017;42(4):222–229.

5. Williams NR, Heifets BD, Blasey C, Sudheimer K, Pannu J, Pankow H, et al. Attenuation of antidepressant effects of ketamine by opioid receptor antagonism. Am J Psychiatry. 2018;175(12):1205–1215.

6. Jelen LA, Lythgoe DJ, Stone JM, Young AH, Mehta MA. Effect of naltrexone pretreatment on ketamine-induced glutamatergic activity and symptoms of depression: a randomized crossover study. Nat Med. 2025;31(9):2958–2966.

7. Bonaventura J, Lam S, Carlton M, Boehm MA, Gomez JL, Solís O, et al. Pharmacological and behavioral divergence of ketamine enantiomers: implications for abuse liability. Mol Psychiatry. 2021;26(11):6704–6722.

8. Jiang Q, Han J, Fine EJ, Ramos-Gonzalez N, et al. Structural basis of opioid receptor activation by PCP and ketamine. Nat Struct Mol Biol. 2026. doi:10.1038/s41594-026-01839-y.

9. Jiang C, DiLeone RJ, Pittenger C, Duman RS. The endogenous opioid system in the medial prefrontal cortex mediates ketamine’s antidepressant-like actions. Transl Psychiatry. 2024;14(1):90.

10. Gomes I, Gupta A, Margolis EB, Fricker LD, Devi LA. Ketamine and major ketamine metabolites function as allosteric modulators of opioid receptors. Mol Pharmacol. 2024;106(5):240–252.

11. Zanos P, Moaddel R, Morris PJ, Georgiou P, Fischell J, Elmer GI, et al. NMDAR inhibition-independent antidepressant actions of ketamine metabolites. Nature. 2016;533(7604):481–486.

12. Zarate CA Jr, Brutsche N, Laje G, Luckenbaugh DA, Venkata SL, Ramamoorthy A, et al. Relationship of ketamine’s plasma metabolites with response, diagnosis, and side effects in major depression. Biol Psychiatry. 2012;72(4):331–338.

13. Gupta A, Devi LA, Gomes I. Potentiation of μ-opioid receptor-mediated signaling by ketamine. J Neurochem. 2011;119(2):294–302.

14. Krupitsky E, Burakov A, Romanova T, Dunaevsky I, Strassman R, Grinenko A. Ketamine psychotherapy for heroin addiction: immediate effects and two-year follow-up. J Subst Abuse Treat. 2002;23(4):273–283.

15. Krupitsky EM, Burakov AM, Dunaevsky IV, Romanova TN, Slavina TY, Grinenko AY. Single versus repeated sessions of ketamine-assisted psychotherapy for people with heroin dependence. J Psychoactive Drugs. 2007;39(1):13–19.

16. McKendrick G, Garrett H, Jones HE, McDevitt DS, Sharma S, Silberman Y, et al. Ketamine blocks morphine-induced conditioned place preference and anxiety-like behaviors in mice. Front Behav Neurosci. 2020;14:75.

17. Zhai H, Wu P, Chen S, Li F, Liu Y, Lu L. Effects of scopolamine and ketamine on reconsolidation of morphine conditioned place preference in rats. Behav Pharmacol. 2008;19(3):211–216.

18. Gupta A, Gomes I, Osman A, Fujita W, Devi LA. Regulation of cannabinoid and opioid receptor levels by endogenous and pharmacological chaperones. J Pharmacol Exp Ther. 2024;391(2):279–288.

19. Gupta A, Décaillot FM, Gomes I, Tkalych O, Heimann AS, Ferro ES, et al. Conformation state-sensitive antibodies to G-protein-coupled receptors. J Biol Chem. 2007;282(8):5116–5124.*

20. Gomes I, Bobeck EN, Margolis EB, Gupta A, Sierra S, Fakira AK, et al. Identification of GPR83 as the receptor for the neuroendocrine peptide PEN. Sci Signal. 2016;9(425):ra43.*

21. Gershman H, Powers E, Levine L, Van Vunakis H. Radioimmunoassay of prostaglandins, angiotensin, digoxin, morphine and adenosine-3’,5’-cyclic-monophosphate with nitrocellulose membranes. Prostaglandins. 1972;1(5):407–423.

22. Gupta A, Gullapalli S, Pan H, Ramos-Ortolaza DL, Hayward MD, Low MJ, et al. Regulation of opioid receptors by their endogenous opioid peptides. Cell Mol Neurobiol. 2021;41(5):1103–1118.*

23. Pan H, Nanno D, Che FY, Zhu X, Salton SR, Steiner DF, et al. Neuropeptide processing profile in mice lacking prohormone convertase-1. Biochemistry. 2005;44(12):4939–4948.

24. Fricker LD, Berman YL, Leiter EH, Devi LA. Carboxypeptidase E activity is deficient in mice with the fat mutation. Effect on peptide processing. J Biol Chem. 1996;271(48):30619–30624.

25. Ellgren M, Spano SM, Hurd YL. Adolescent cannabis exposure alters opiate intake and opioid limbic neuronal populations in adult rats. Neuropsychopharmacology. 2007;32(3):607–615.

26. Spano MS, Ellgren M, Wang X, Hurd YL. Prenatal cannabis exposure increases heroin seeking with allostatic changes in limbic enkephalin systems in adulthood. Biol Psychiatry. 2007;61(4):554–563.

27. Shen H, Moussawi K, Zhou W, Toda S, Bhatt DK, Bhatt S, et al. Heroin relapse requires long-term potentiation-like plasticity mediated by NMDA2b-containing receptors. Proc Natl Acad Sci U S A. 2011;108(48):19407–19412.

28. Munguba H, Arefin A, Hasegawa R, Posa L, Romano GR, Peddada TN, et al. Mechanism-guided identification of antidepressant G protein-coupled receptor drug targets. Cell. 2026;189(9):2612–2632.e24.

29. Fricker LD, Osman A, Gupta A, Gomes I, Devi LA. Antidepressants and the endogenous opioid system. Biochem Pharmacol. 2025;242(Pt 4):117392.

30. Williams JT, Christie MJ, Manzoni O. Cellular and synaptic adaptations mediating opioid dependence. Physiol Rev. 2001;81(1):299–343.

31. Drakenberg K, Nikoshkov A, Horvath MC, Fagergren P, Gharibyan A, Saarelainen K, et al. Mu opioid receptor A118G polymorphism in association with striatal opioid neuropeptide gene expression in heroin abusers. Proc Natl Acad Sci U S A. 2006;103(20):7883–7888.

32. Heimer L, Zahm DS, Churchill L, Kalivas PW, Wohltmann C. Specificity in the projection patterns of accumbal core and shell in the rat. Neuroscience. 1991;41(1):89–125.

33. Marchant NJ, Millan EZ, McNally GP. The hypothalamus and the neurobiology of drug seeking. Cell Mol Life Sci. 2012;69(4):581–597.

34. Yu J, Ishikawa M, Wang J, Schlüter OM, Sesack SR, Dong Y. Ventral tegmental area projection regulates glutamatergic transmission in nucleus accumbens. Sci Rep. 2019;9(1):18451.

35. Bossert JM, Stern AL, Theberge FR, Marchant NJ, Wang HL, Morales M, et al. Role of projections from ventral medial prefrontal cortex to nucleus accumbens shell in context-induced reinstatement of heroin seeking. J Neurosci. 2012;32(14):4982–4991.

36. Dalton GL, Wu DC, Wang YT, Floresco SB, Phillips AG. NMDA GluN2A and GluN2B receptors play separate roles in the induction of LTP and LTD in the amygdala and in the acquisition and extinction of conditioned fear. Neuropharmacology. 2012;62(2):797–806.

37. Saland SK, Kabbaj M. Sex differences in the pharmacokinetics of low-dose ketamine in plasma and brain of male and female rats. J Pharmacol Exp Ther. 2018;367(3):393–404.

