## Supplemental Information for "Ketamine Modulates Endogenous Opioid Peptide Signaling and Reduces Heroin-Seeking in Male Long-Evans Rats"

##### **Supplemental Methods:**

###### *Quantitative real-time PCR*

ThermoFisher Taqman probes used: Oprm1 (Rn01430371\_m1), Oprk1 (Rn01448892\_m1), Pdyn (Rn00571351\_m1), Penk (Rn00567566\_m1), Grin2a (Rn00561341\_m1), and Grin2b (Rn00680474\_m1). Gapdh (Rn01775763\_g1) Genes of interest were run on the FAM channel, and Gapdh was run on the VIC channel of a Roche LightCycler II.

###### *Membrane preparation*

20 mM Tris-Cl buffer used to homogenize tissue contained 250 mM sucrose, 2 mM EGTA, and 1 mM MgCl<sub>2</sub> (pH 7.4). 2mM Tris-Cl buffer used to resuspend pellet contained 2 mM EGTA and 10% glycerol (pH 7.4).

###### ELISA

*o*-phenylenediamine (5 mg *o*-phenylenediamine/10 ml of citric phosphate buffer; 0.15 M citric acid + 0.15 M dibasic sodium phosphate, pH 5.0 containing 20 μl hydrogen peroxide). 5G8 antibody concentration (1:500 in PBS containing 1% BSA). Anti-mouse IgG coupled to horseradish peroxidase concentration (1:1000 in PBS containing 1% BSA)

Commented [DL1]: Need more details - amount of membrane protein, dil of abs, incubation time etc. @lvone - please fill it in.

Commented [IG1R2]: Done

###### [<sup>35</sup>S] GTPγS binding

Assay buffer used contains (50 mM Tris-Cl, pH 7.4, with 100 mM NaCl, 10 mM MgCl<sub>2</sub>, 0.2 mM EGTA, and protease inhibitor cocktail). Filters used with Brandal filtration system are GF/B filters.

Commented [DL2]: Need more details -time of inc. buffer used etc. @lvone - please fill it in.

Commented [IG2R2]: Done

###### *Radioimmunoassay*

After acid extraction overnight samples were stored at -70°C. 50 mM sodium phosphate buffer buffer used to resuspend samples contained 0.1% Triton X-100 and

0.1% BSA. Assay buffer A consisted of 50 mM sodium phosphate buffer, pH 7.6 containing 0.1% Triton X-100 and 0.1% BSA. Range of Leu-enk standards used (1pM – 10 μM), rabbit anti-Leu-enkephalin IgG antibody (1:1000 in assay buffer A) and [$^{125}$I]-Leu-enkephalin (20,000 cpm/100μl/tube).

#### **Heroin Self-Administration Cohort**

##### *Animals and Surgical Procedures*

Post-surgical care included a three-day recovery period during which catheters were flushed daily with heparinized saline containing ampicillin. Catheter patency was confirmed using Brevital prior to experimentation.

##### *Intravenous Heroin Self-Administration*

Catheters were flushed with heparinized saline at the end of each session.

##### *Forced Abstinence and Ketamine Interventions*

The intravenous dose was selected to approximate plasma concentrations achieved with the most validated route of administration.

### **Supplementary Figure Legends**

**Supplementary Figure 1. Acute ketamine does not alter endogenous opioid system gene expression in the PFC or dorsal striatum.** (A) *Oprm1*, *Oprk1*, *Pdyn*, and *Penk* mRNA levels in the PFC following saline, 3 mg/kg, or 10 mg/kg ketamine at 1, 3, and 7 days post-injection. (B) *Oprm1*, *Oprk1*, *Pdyn*, and *Penk* mRNA levels in the dorsal striatum. No significant changes were observed in either region. Data are mean ± SEM.

**Supplementary Figure 2. Acute ketamine does not alter glutamatergic gene expression in Cohort 1.** (A) *Grin2a* and *Grin2b* mRNA levels in the NAc and PFC following saline, 3 mg/kg, or 10 mg/kg ketamine at 1, 3, and 7 days post-injection. (B) *Grin2a* and *Grin2b* mRNA levels in the dorsal striatum. No significant changes were observed. Data are mean ± SEM.

**Supplementary Figure 3. ELISA and GTPγS binding in whole NAc from Cohort 1.** (A) ELISA revealed elevated MOPr protein levels in the NAc of ketamine-treated animals at the 7-day timepoint (\*\*p<0.01). (B) Raw [$^{35}$S]GTPγS binding (counts per million) in the NAc showing a main effect of treatment (\*\*\*\*p<0.0001). Data are mean ± SEM.

**Supplementary Figure 4. Ketamine effects on the second cue-induced drug-seeking session.** (A) Active and inactive lever presses during the second cue-induced drug-seeking session, 3 days following 10 mg/kg ip ketamine. Ketamine-treated animals pressed the inactive lever significantly fewer times than saline-treated animals (\*\*$p < 0.01$). Data are mean ± SEM.

**Supplementary Figure 5. No significant transcriptional alterations in the NAcC following heroin self-administration and ketamine treatment.** (A) *Oprm1*, (B) *Oprk1*, (C) *Pdyn*, (D) *Penk*, and (E) *Grin2b* mRNA levels in the NAcC of saline- and ketamine-treated animals following the heroin-primed seeking session. No significant differences were observed. Data are mean ± SEM.

**Supplementary Figure 6. No significant transcriptional alterations in the NAcS following heroin self-administration and ketamine treatment.** (A) *Oprm1*, (B) *Oprk1*, (C) *Pdyn*, (D) *Penk*, (E) *Grin2a*, and (F) *Grin2b* mRNA levels in the NAcS of saline- and ketamine-treated animals following the heroin-primed seeking session. No significant differences were observed. Data are mean ± SEM.

**Supplementary Figure 7. No significant transcriptional alterations in the PFc following heroin self-administration and ketamine treatment.** (A) *Oprm1*, (B) *Oprk1*, (C) *Pdyn*, (D) *Penk*, (E) *Grin2a*, and (F) *Grin2b* mRNA levels in the PFc of saline- and ketamine-treated animals following the heroin-primed seeking session. No significant differences were observed. Data are mean ± SEM.

**Supplementary Figure 8. Pearson correlation analyses between heroin self-administration behavior and gene expression.** Correlation matrices between average active lever presses and infusions over the last 3 days of heroin self-administration and qPCR data in (A) NAcC of saline-treated animals, (B) NAcC of ketamine-treated animals, (C) NAcS of saline-treated animals, and (D) NAcS of ketamine-treated animals. Significant positive correlations were observed between heroin-taking behavior and *Pdyn* ($r = 0.73$, $p = 0.012$) and *Penk* ($r = 0.69$, $p = 0.01$) expression in the NAcS of ketamine-treated animals only. \*$p < 0.05$, \*\*$p < 0.01$, \*\*\*$p < 0.001$.
